# Poliovirus-nonsusceptible Vero cell line for the World Health Organization global action plan

**DOI:** 10.1101/2020.08.19.257204

**Authors:** Yuko Okemoto-Nakamura, Kenji Someya, Toshiyuki Yamaji, Kyoko Saito, Makoto Takeda, Kentaro Hanada

**Affiliations:** Department of Biochemistry and Cell Biology, National Institute of Infectious Diseases, 1-23-1 Toyama, Shinjuku-ku, Tokyo, 162-9640, Japan; Department of Virology 3, National Institute of Infectious Diseases, 4-7-1, Gakuen, Musashimurayama, Tokyo, 208-0011, Japan

## Abstract

Polio or poliomyelitis is a disabling and life-threatening disease caused by poliovirus (PV). As a consequence of global polio vaccination efforts, wild PV serotype 2 has been eradicated, and wild PV serotypes 1- and 3-transmitted cases have been largely eliminated except for in limited regions around the world. However, vaccine-derived PV, pathogenically reverted live PV vaccine strains in vaccinated humans, has become a serious issue. For the global eradication of polio, the World Health Organization is conducting the third edition of the Global Action Plan, which is requesting stringent control of potentially PV-infected materials. To facilitate the mission, we generated a PV-nonsusceptible Vero cell subline, which may serve as an ideal replacement of standard Vero cells to isolate emerging/re-emerging viruses without the risk of generating PV-infected materials.

## Introduction

Polio or poliomyelitis is a disabling and life-threatening disease caused by poliovirus (PV), a nonenveloped, positive-sense single-stranded RNA virus classified in the genus *Enterovirus* in the family *Picornaviridae* ^1^, of which there are three wild serotypes: PV 1–3. As a consequence of global polio vaccination efforts, wild PV serotype 2 has been eradicated, and wild PV serotypes 1- and 3-transmitted cases have been largely eliminated except for in limited regions around the world. However, vaccine-derived PV (VDPV), pathogenically reverted live PV vaccine strains in vaccinated humans, has become a serious issue. In particular, serotype 2 oral polio vaccine (OPV) has been the predominant cause of VDPV cases since the eradication of wild PV serotype 2 in 1999 ^2, 3^.

For the global eradication of polio, the World Health Organization (WHO) is conducting the third edition of the Global Action Plan (GAPIII) ^4^. The WHO GAPIII rationalizes that with the use of inactivated PV vaccines, most countries will have no need to retain live PV in the post-eradication and post-OPV era, and that facility-associated risks can be eliminated by destruction of all wild PV and OPV infectious and potentially infectious materials. However, when PV-susceptible cells are used to isolate non-PV viruses for public health purposes in countries or regions with wild PV- and/or OPV-derived cases, the resultant cell culture has to be handled as potentially infectious, making it a costly and laborious burden.

The Vero cell line is a continuous cell line established from the kidney of the *Chlorocebus sabaeus* species (or the *sabaeus* subspecies of *C. aethiops*) of African green monkeys (AGMs) ^5, 6^, the whole genome sequences of which have been determined in several sublines ^6, 7^. Vero cells are highly susceptible to various viruses, including PV, in part because of the loss of the type I interferon gene cluster ^6^, and they have been successfully used as a first-choice cell model for various types of life-threatening emerging pathogens ^8 9–12^. Vero cells are also frequently used as a cell substrate for human virus vaccines because of their non-tumorigenicity ^13–15^.

To provide an ideal replacement of standard Vero cells to isolate emerging/re-emerging viruses without the risk of generating potentially PV-infected materials, we here generated a PV-nonsusceptible Vero cell subline by disrupting genomic genes encoding AGM paralogs of the human PV receptors.

## Results and Discussion

### Genomic genes and cDNAs of the human PV receptor homologs in Vero cells

The cell receptor of PV (PVR; also known as CD155) is a glycosylated membrane protein with an *N*-terminal extracellular domain, single transmembrane domain, and *C*-terminal cytoplasmic domain (Fig. 1a) ^1^. The human genome has only one PVR-encoding gene, which expresses four alternative splicing transcripts. Among the transcript variants, only two variants have the transmembrane domain and function as the cell receptor of PV ^1^. The AGM genome contains two paralogs of the human *PVR* ^16, 17^, hereafter referred to as *PVR1* and *PVR2*, which probably arise because of gene duplication. A previous study showed that both AGM PVRs have the potential to form functional PVRs ^17^.

**Figure 1.**
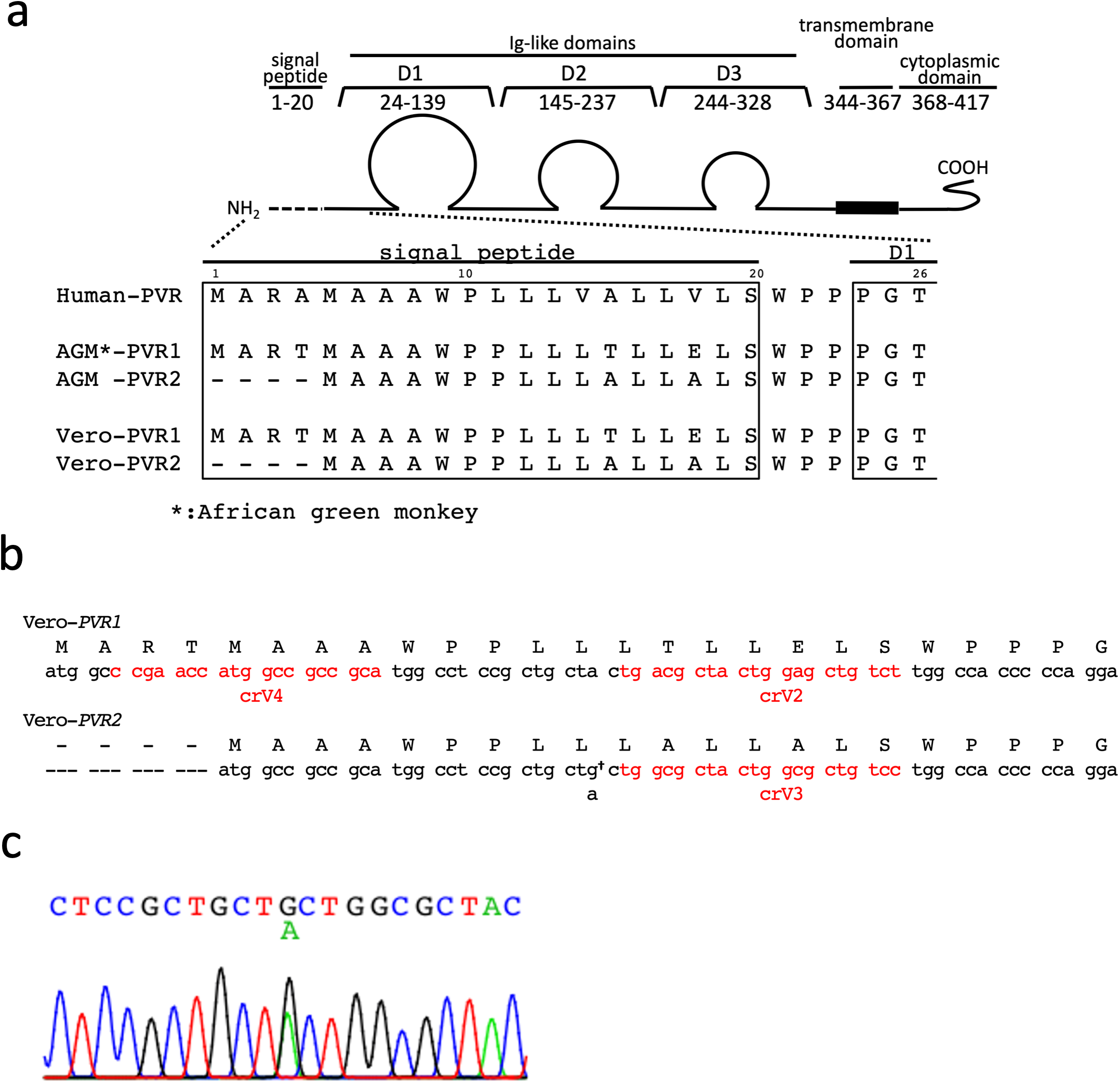
Generation of *PVR1*/*PVR2* DKO Vero cell lines. (**a)** Diagram of PVR. The signal peptide and the N-terminus of the first domain (D1) of the three Ig-like domains, which is responsible for viral recognition, is encoded by exon 1 of the human *PVR* gene. Amino acid sequences are aligned with AGM (*Chlorocebus sabaeus*) or Vero *PVR*s (see also Supplementary Table 1). (**b**) Nucleotide sequences of the initial ATG codon and downstream region in Vero-*PVR1* and Vero-*PVR2.* Genomic DNA of Vero cells were extracted and the coding region located on exon 1 of *PVR1* and *PVR2* were sequenced. DNA sequences of Vero-*PVR1* and Vero-*PVR2* are shown in lowercase letters; the predicted amino acid sequences are shown in capital letters. Vero-*PVR2* was found to have a single-nucleotide variation (SNV) (†) in the open reading frame (ORF), probably due to heterozygosity of the gene. (**c**) DNA sequencing chromatogram of a Vero-*PVR2* region. Sanger sequencing of Vero-*PVR2* showed an SNV in the ORF of exon 1 (see also *panel b*).

In both genes, the first exon encodes the translational initiation codon, the signal peptide domain, and a part of the extracellular immunoglobulin-like V-type domain, which are essential for functional PVR ^1, 17^. We amplified the first exon and flanking introns of the PVRs in the Vero JCRB9013 cell line (which is equivalent to the Vero ATCC CCL-81 cell line) by genomic polymerase chain reaction (PCR), and sequenced them to design guide RNA (gRNA) targeting this exon. The amplified region of the *PVR2* gene of Vero cells was found to contain synonymous single nucleotide variations (SNVs) when compared with counterparts in AGM reference sequences (Fig. 1b, c). Additionally, we obtained several full-length cDNA clones of Vero cell *PVR1* and *PVR2* and sequenced them individually. Whereas only one type of sequence was obtained from the *PVR1* cDNA set (Supplementary Fig. 1a), two types were obtained from the *PVR2* cDNA set (Supplementary Fig. 1b), which indicates that different patterns of SNVs, including synonymous and nonsynonymous mutations, exist between the two alleles of *PVR2* gene. Of note, Vero cells are pseudodiploid ^6, 7^.

### Disruption of *PVR1*, *PVR2*, or both in Vero cells

Following the scheme shown in Fig. 2a, we constructed *PVR*-disrupted Vero cell lines, obtaining two *PVR1* single-knockouts (SKOs), two *PVR2* SKOs, and six *PVR1*/*PVR2* double-knockouts (DKOs) cell clones (Supplementary Fig. 2). Among the *PVR1*/*PVR2* DKO cell clones, two clones (Δ*PVR1/2*-1 and Δ*PVR1/2*-2) were derived from a *PVR1* SKO clone (Δ*PVR1*-1) while four clones (Δ*PVR1/2*-3, Δ*PVR1/2*-4, Δ*PVR1/2*-5, and Δ*PVR1/2*-6) were derived from another *PVR1* SKO clone (Δ*PVR1*-2; Supplementary Fig. 2). No obvious morphological changes were observed among these clones (Fig. 3a). Nevertheless, the Δ*PVR1*-2 cell clone had retarded entry into the logarithmic phase of cell growth after plating, when compared with the parental, Δ*PVR1*-1, Δ*PVR2*-1, and Δ*PVR2*-2 cell clones (Fig. 3b). Thus, we chose Δ*PVR1*-1, not Δ*PVR1*-2, as the Vero Δ*PVR1* representative for depositing to a cell bank (see also below).

**Figure 2.**
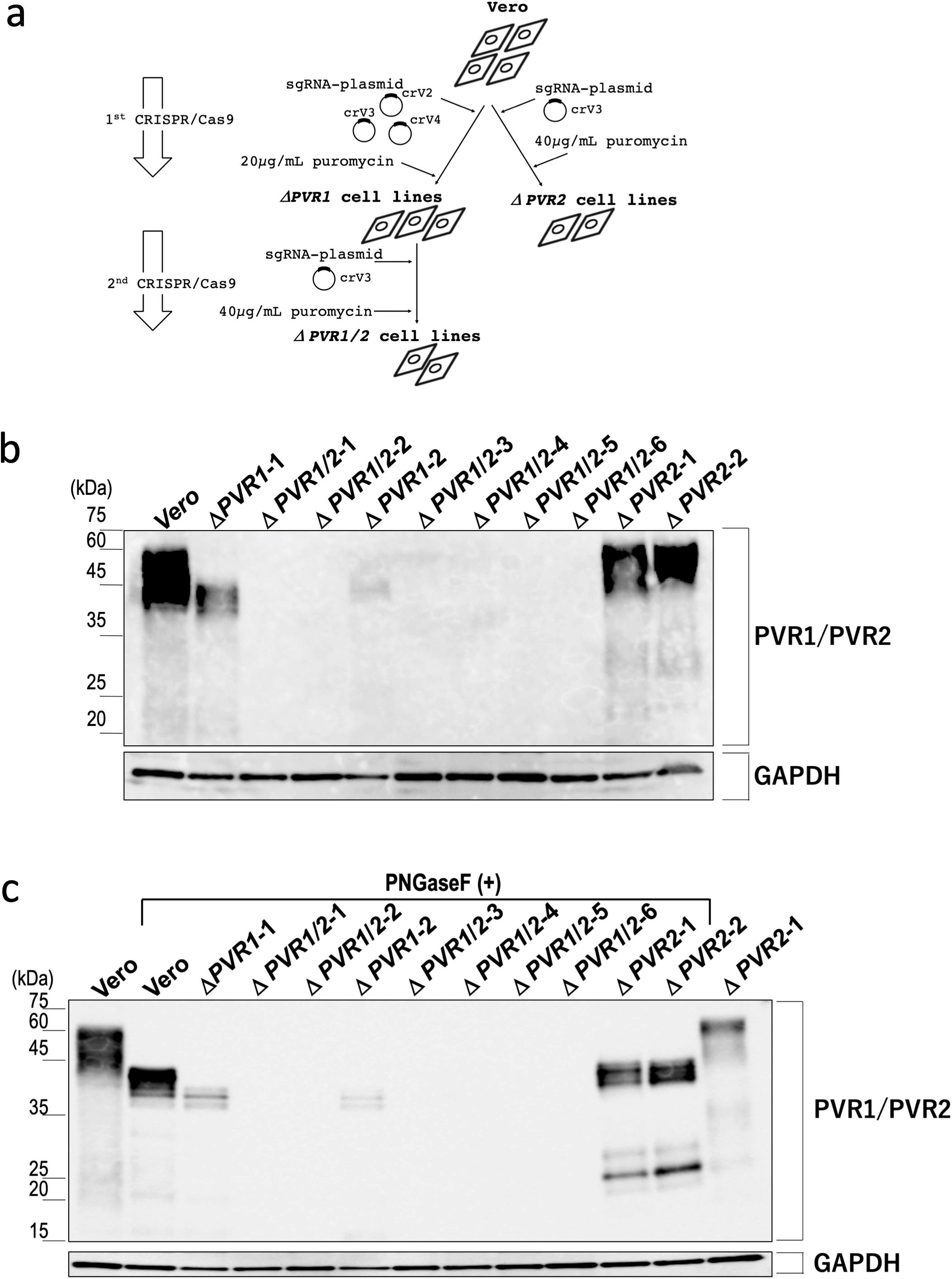
Generation of *PVR1*/*PVR2* double-knockout (DKO) cell lines. (**a**) Diagram for construction of *PVR1*/*PVR2* double-deficient cell lines. In the first-round experiment, co-transfection of parental Vero cells with two plasmids expressing the sgRNA crV4 (targeting to *PVR1*) or crV3 (targeting to *PVR2*) gave only *PVR1* single-knockout (SKO) cell lines as shown (Supplementary Fig. 2). One of the purified cell clones named Vero Δ*PVR1*-1 was subjected to a second-round experiment, in which the cells were transfected with the crV3-expressing plasmid, and two *PVR1/2* DKO cell lines named Vero Δ*PVR1/2*-1 and Vero Δ*PVR1/2*-2, respectively, were obtained (see also Supplementary Fig. 2). In another experiment, transfection of Vero cells with the plasmid expressing crV2 (targeting to *PVR1*) gave a *PVR1* SKO cell line named Vero Δ*PVR1*-2, and transfection of Vero Δ*PVR1*-2 cells with the crV3-expressing plasmid gave four *PVR1/2* DKO cell lines: Vero Δ*PVR1/2*-3, Vero Δ*PVR1/2*-4, Vero Δ*PVR1/2*-5, and Vero Δ*PVR1/2*-6 (Supplementary Fig. 2). For construction of *PVR2* SKO cells, Vero JCRB9013 cells were transfected with the crV3-expressing plasmid, and two mutant cell lines, Vero Δ*PVR2*-1 and Vero Δ*PVR2*-2, were obtained (see also Supplementary Fig. 2). (**b**) Fifteen-microgram aliquots of protein samples prepared from the Vero cell line (Vero), two *PVR1* SKO cell lines (Δ*PVR1*-1 and Δ*PVR1*-2), six *PVR1*/*PVR2* DKO cell lines (Δ*PVR1/2*-1, Δ*PVR1/2*-2, Δ*PVR1/2*-3, Δ*PVR1/2*-4, Δ*PVR1/2*-5, and Δ*PVR1/2*-6), and two *PVR2* SKO cell lines (Δ*PVR2*-1 and Δ*PVR2*-2) were subjected to western blot analysis using the anti-PVR/CD155 rabbit monoclonal antibody D3G7H (upper panel). Loading control GAPDH was detected with the anti-GAPDH rabbit monoclonal antibody 14C1 (lower panel). (**c**) After removal of N-linked glycans, protein samples equivalent to 5 μg of cell lysate were subjected to western blot analysis. PNGase F-untreated controls of Vero and Δ*PVR2*-1 cell-derived samples are shown in the leftmost and rightmost lanes, respectively, in the panel.

**Figure 3.**
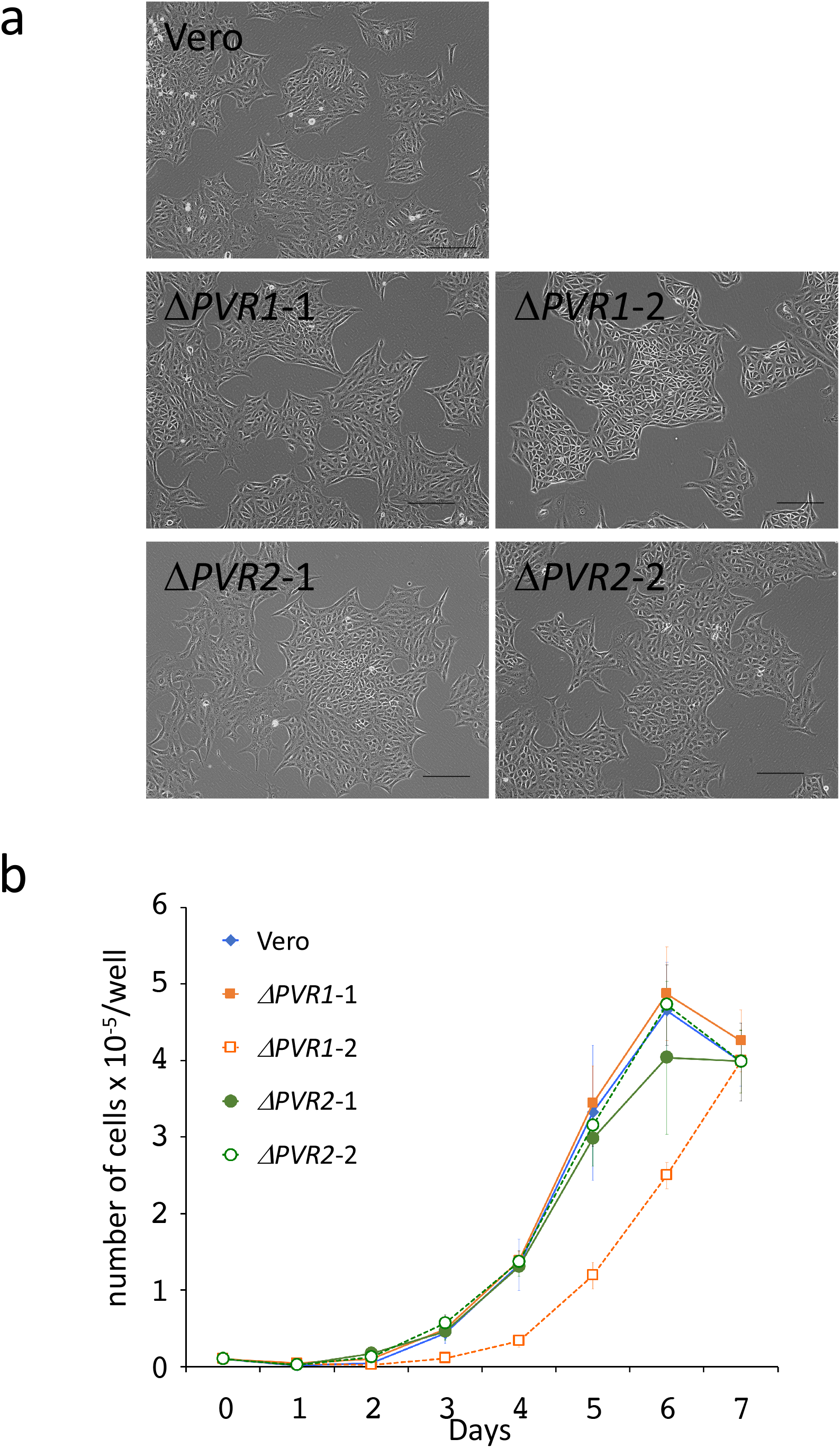
Cell morphology and growth curve of Vero and *PVR* knockout cell lines. (**a**) Cell morphology of the Vero cell line, two independent *PVR1* SKO cell lines (Δ*PVR1*-1 and Δ*PVR1*-2), and two independent *PVR2* SKO cell lines (Δ*PVR2*-1 and Δ*PVR2*-2; see also Supplementary Fig. 2). Scale bar: 200 μm. (**b**) Cells were seeded into 24-well plates at 1 × 10^4^ cells/well and cultured for seven days. Live cells were counted each day and the proliferation curves are shown (means ± Standard Deviation, *n* = 3).

In western blot analysis with an anti-human PVR monoclonal antibody, which was expected to cross-react with AGM PVRs, parental Vero cells exhibited broad bands, with band mobility corresponding to ~60–40 kDa (Fig. 2b). As peptide-*N*-glycosidase F (PNGase F)-treatment sharpened the bands (Fig. 2c), the observed band broadness was largely due to the heterogeneity of protein glycosylation. Compared with the parental control, the band signal was greatly reduced with Vero *PVR1* SKO cells, but not with Vero *PVR2* SKO cells. The signal retained with Vero *PVR1* SKO cells disappeared with Vero *PVR1*/*2* DKO cells, which indicates that the band represented the PVR2 protein (Fig. 2b, c).

### PV-nonsusceptible Vero cell lines

Infectivity of PV in these Vero cell clones was analyzed. The PV Sabin serotypes 1 and 3 strains underwent multiple rounds of infection very efficiently in parental Vero cells, while no virus production was observed in Vero Δ*PVR1*/*2* cells (Fig. 4a–d, Supplementary Fig. 3e, f). Even when cells were infected with PVs at a high multiplicity of infection (MOI) of 10, neither virus production nor viral RNA synthesis was detected (Fig. 4e–h, Supplementary Fig. 3g–j). Notably, *PVR1* SKO was enough to abrogate PV susceptibility (Fig. 4e–h, Supplementary Fig. 3a, b), while *PVR2* SKO had no discernible impact (Fig. 4e–h, Supplementary Fig. 3c, d, g–j), which indicates that PVR1 is the predominant receptor for PV infection in Vero cells. It is currently unknown why the endogenous *PVR2* in Vero cells does not seem to serve as a functional PVR, although ectopic overexpression of AGM *PVR2* in mouse L cells endows PV-susceptibility to the cells ^17^.

**Figure 4.**
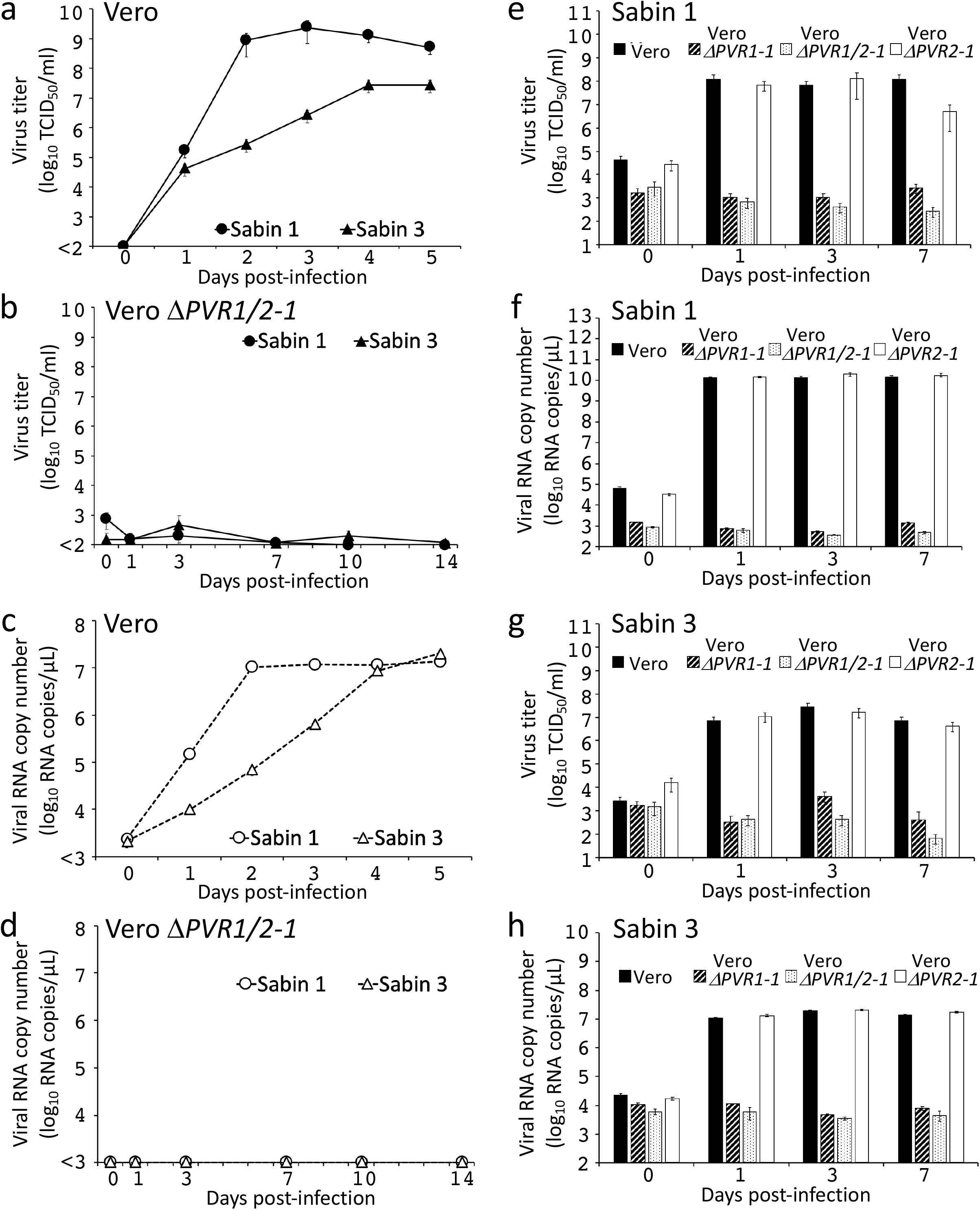
PV replication and viral RNA synthesis in the parental and *PVR*-disrupted Vero cell lines. (**a**–**d**) The indicated cell lines were infected with PV Sabin 1 and 3 strains at MOI of 0.01 (**a**–**d**) and 10 (**e**–**h**), and virus titers (**a, b, e, g**) and viral RNA (**c, d, f, h**) in the culture fluids were quantified at the indicated times post-infection (mean ± standard deviation, *n* = 3). See also Supplementary Fig. 3 for other Vero *PVR*-disrupted cell clones established in this study.

### Susceptibility to non-PV viruses

As expected, all the *PVR1*- and/or *PVR2*-disrupted Vero cell clones remained susceptible to measles virus (MV; AIK-C vaccine strain-derived recombinant virus; Supplementary Fig. 4), rubella virus (RV; rHS717AG1 wild strain-derived recombinant virus; Supplementary Fig. 5), and Japanese encephalitis virus (JEV; Nakayama strain; Supplementary Fig. 6), eliminating the possibility of non-specific virus tolerance of these Vero cell clones.

Compared with parental Vero cells, all *PVR* KO clones retained high susceptibility to MV and RV, with similar time courses in virus production (Supplementary Fig. 4, 5). All *PVR* KO clones also retained high susceptibility to JEV. However, the JEV tilter levels of Vero, Δ*PVR1*-1, and Δ*PVR1*-1-derived *PVR1/2* DKO clones, and *PVR2* SKO clones peaked at three days post-infection (Supplementary Fig. 6b, c) while those of Δ*PVR1*-2 and Δ*PVR1*-2-derived *PVR1/2* DKO cell clones (except for Δ*PVR1/2*-6) peaked two days post-infection (Supplementary Fig. 6d). The cause of the different characteristics of Vero Δ*PVR1*-1 and Δ*PVR1*-2 clones is unknown.

Among the Vero cell mutant clones established in this study, we chose Vero Δ*PVR1*-1, Vero Δ*PVR2*-1, and Vero Δ*PVR1/2*-1 clones as representative cell lines of Vero *PVR1* SKO, Vero *PVR2* SKO, and Vero *PVR1*/*2* DKO cell clones, respectively (Supplementary Fig. 2). It should be noted that the phenotypes of the representative cell lines were not due to any clonal effects as other mutant cell lines established in this study broadly exhibited the same phenotypes (Figs. 2–4 and Supplementary Figs. 3–6). For the global distribution, these representative cell lines have been deposited in the Japanese Collection of Research Bioresources (JCRB) Cell Bank, a non-profit public cell bank in Japan. Of note, the representative cell lines deposited to the cell bank were renamed Vero Δ*PVR1,* Vero Δ*PVR2,* and Vero Δ*PVR1/2,* respectively, for simplicity (Supplementary Fig. 2).

When an outbreak of an emerging and re-emerging infectious disease occurs, the identification of the causative pathogen is crucial. Additionally, in vitro pathogen culture systems are invaluable for fully characterizing the pathogen and the development of anti-pathogenic drugs. The present study provides evidence that when Vero Δ*PVR1/2* cells are employed as host cells for non-PV viruses, the resultant cell culture can be excluded from classification as “potentially PV-infected materials”. Thus, Vero Δ*PVR1/2* cells may serve as an ideal replacement of standard Vero cells to isolate emerging/re-emerging viruses without the risk of generating potentially PV-infected materials, which is in alignment with the WHO GAPIII mission.

## Methods

### Cells and cell culture

Vero cells (JCRB9013) were obtained from the Japanese Collection of Research Bioresources (JCRB) Cell Bank. Note that the quality of the cells was assured by standard tests of the cell bank. Cells were cultivated in a normal growth medium consisting of Eagle’s minimal essential medium (MEM; Fujifilm Wako Pure Chemical Co., Osaka, Japan) supplemented with 5% heat-inactivated fetal bovine serum (FBS; Sigma-Aldrich, MO, USA) and penicillin-streptomycin (Fujifilm Wako Pure Chemical Co.), at 37°C and under an atmosphere of 5% CO_2_ and 100% humidity. Cells were detached with AccuMax dissociation solution (Innovative Cell Technologies, CA, USA) and passaged at a split ratio of 1:5.

### Cell growth curve and microscopy

Cells were seeded at a density of 1 × 10^4^ cells/well in a 24-well plate (Corning Inc, N.Y., USA) and cultured for seven days in normal growth medium. Cells were detached with AccuMax and stained with trypan blue solution (Thermo Fisher Scientific), and live cells in three independent wells were counted using a Countess II Automated Cell Counter (Thermo Fisher Scientific). To examine cell morphologies, cells were photographed using a BZ-X710 All-in-One Fluorescence Microscope (Keyence, Osaka, Japan).

### cDNA sequencing of Vero-*PVR1* and Vero-*PVR2*

RNA was extracted from Vero cells using the NucleoSpin RNA kit (Takara Bio Inc.), and cDNA was synthesized with the PrimeScript 1st strand cDNA Synthesis Kit (Takara Bio Inc.). Full-length Vero-*PVR1* and Vero-*PVR2* cDNA was amplified with the Easy-A High Fidelity PCR Cloning Enzyme (Agilent Technologies, CA, USA), and cloned into T-vector pMD20 (Takara Bio Inc.). To amplify Vero-*PVR1*, cloning primers 1-Fw; 5′-ATG GCC CGA ACC ATG GCC GCC GCA TG-3′ and 1/2-Rv; 5′-CCT TGT GCC CTC TGT CTG TGG ATC CTG-3′ were used. For Vero-*PVR2*, two primer sets were designed because our genomic DNA sequence analysis showed that *PVR2* has a nucleotide polymorphism at its 5′ terminus (Supplementary Fig. 1b, c). One of the primer sets for Vero-*PVR2* was cloning primer 2G-Fw (5′-ATG GCC GCC GCA TGG CCT CCG CTG CTG-3′) and 1/2-Rv (5′-CCT TGT GCC CTC TGT CTG TGG ATC CTG-3′), and the other primer set was 2A-Fw (5′-ATG GCC GCC GCA TGG CCT CCG CTG CTA-3′) and 1/2-Rv (5′-CCT TGT GCC CTC TGT CTG TGG ATC CTG-3′). The nucleotide sequences were determined by Sanger sequencing.

### DNA sequencing of Vero-*PVR1* and Vero-*PVR2*

Genomic DNA was isolated from Vero cells using the NucleoSpin Tissue kit (Takara Bio Inc., Shiga, Japan). To determine the sequence of the N-terminal *PVR1*- and *PVR2*-coding regions, the genomic DNA served as a template for PCR with Tks Gflex DNA polymerase (Takara Bio Inc.) using the following primer sets: primer set-1 for *PVR1* (primer 1-Fw; 5′-ATC TGT CCC ATC ACG AGT TGA-3′, primer 1-Rv; 5′-AAC CCC GGG GCA GAG ATG-3′), and primer set-2 for *PVR2* (primer 2-Fw; 5′-CTC GGC TCA CTG GTC TGG AAC-3′, primer 2-Rv; 5′-CTG CAT GCG GAG CTG GGA GC-3′). The resultant PCR products were subjected to Sanger DNA sequencing.

### Construction of a vector for the CRISPR/Cas9 system

Guide sequences containing the target sequences were cloned into the BsmBI site of the pSELECT-CRISPR cas9 plasmid as described previously ^18, 19^. The target sequences specific for Vero-*PVR1* and -*PVR2* are shown in Fig. 1b. Vero cells were transfected with the constructed plasmids using the Lipofectamine 2000 reagent (Thermo Fisher Scientific, MA, USA) according to the manufacturer’s protocol. Three days after transfection, cells were cultivated for three days in growth medium supplemented with 20 μg/ml of puromycin (Takara Bio Inc.) for *PVR1* KO or 40 μg/ml for *PVR2* KO. Puromycin-resistant colonies were further grown without puromycin, isolated using cloning cups, and subcultured.

### Western blotting

All procedures were performed at 4° C or on ice, unless otherwise noted. Cells were washed with phosphate buffered saline (PBS) three times, removed from the plates using cell scrapers (IWAKI, Tokyo, Japan) in 1 ml of PBS, and centrifuged at 10,000 × *g* for 3 min. The cell pellets were stored at −30°C until analysis. Each frozen pellet was thawed on ice and lysed for 10 min with 200 μl of 100 mM Tris-HCl (pH 7.0) buffer containing 0.5% NP-40, 0.5% sodium deoxycholate, 10 mM ethylenediaminetetraacetic acid (EDTA), and 100 mM NaCl. After centrifugation at 1,000 × *g* for 3 min to remove insoluble debris, the supernatant fraction was collected as cell lysate. Protein concentrations of cell lysates were measured using a bicinchoninic acid protein assay kit (Takara Bio Inc.). Cell lysate containing the same amount of protein was mixed with 10 volumes of methanol and then centrifuged at 20,000 × *g* for 30 min to precipitate proteins. After drying by air, the precipitated proteins were dissolved and denatured in modified Laemmli sample buffer (Bio-Rad, CA, USA), containing100 mM dithiothreitol rather than 5% 2-mercaptoethanol, by boiling for 10 min. Fifteen micrograms of protein was electrophoresed on 15% e-PAGEL sodium dodecyl sulfate polyacrylamide gel electrophoresis (SDS-PAGE) gels (ATTO, Tokyo, Japan) and transferred to polyvinylidene difluoride membranes (ATTO). The anti-PVR/CD155 rabbit monoclonal antibody D3G7H (Cell Signaling Technology, MA, USA; 0.1 μg/ml) and anti-GAPDH rabbit monoclonal antibody 14C10 (Cell Signaling Technology; 0.1 μg/ml) were diluted with Can Get Signal Immunoreaction Enhancer Solution (TOYOBO Life Science, Osaka, Japan) and used as primary antibodies. A horseradish peroxidase (HRP)-labeled anti-rabbit IgG antibody (Jackson ImmunoResearch, PA, USA; 0.7 μg/ml) in Tris-buffered saline with 0.05% Tween 20 was used as a secondary antibody. Proteins were detected by Immobilon Western Chemiluminescent HRP Substrate (Merck Millipore, MA, USA) and a LAS-3000 mini chemiluminescence imaging system (Fujifilm, Tokyo, Japan) with ImageGauge software (Fujifilm).

### Deglycosylation of proteins

To remove N-linked glycans, 25 μg of protein was precipitated from cell lysate as described in Methods and treated with peptide-N-glycosidase F (PNGase F; New England Biolabs, MT, USA) according to the manufacturer’s instructions.

### PV infection and titration

Vero and *PVR* KO cells seeded in a 12-well plate were infected with PV Sabin 1 and Sabin 3 strains (National Institute for Biological Standards and Control, Hertfordshire, UK) at MOI of 10 or 0.01. After 1 h incubation at 35°C, the cells were washed three times with PBS (−) and cultured at 35°C in a 5% CO_2_ humid atmosphere with MEM (Thermo Fisher Scientific) containing 2% FBS (Biowest, Nuaillé, France) and penicillin-streptomycin (Cosmo Bio Co., Tokyo, Japan). At appropriate intervals, cells were dissociated by pipetting and collected together with culture supernatants as the culture fluid. The culture fluid was subjected to three freeze-thaw cycles and then centrifuged at 1,000 × *g* for 5 min to remove debris. Virus titer was determined by 50% end-point dilution assay (50% cell culture infectious dose, TCID_50_) using HEp-2 cells (ATCC CCL-23), a HeLa sub-line (ATCC, VA, USA). Viral RNA copy numbers in the culture fluid was also measured by real-time reverse transcription-PCR ^20, 21^.

## Acknowledgments

We are grateful to Drs. Satoshi Koike and Hiroyuki Shimizu for their expertise in poliovirus research and invaluable suggestions. We thank Martin Cheung PhD, from Edanz Group (https://en-author-services.edanzgroup.com/ac) for editing a draft of this manuscript.

## Author contributions

K.H. and M.T. designed research studies. Y.O.-N. and T.Y. designed sgRNAs. Y.O.-N. isolated PVR-disrupted cell lines and performed cell experiments. Ky.S and Y.O.-N. performed JEV experiments, and Ke.S. performed all other viral experiments. All authors interpreted the data. K.H., Y.O.-N., and Ke.S. drafted the manuscript and all authors approved the final version of manuscript.

## Competing interests

We declare that we have no conflicts of interest.

## Funding

This research was supported by AMED-CREST (JP19gm0910005j0005 to K.H.), JSPS KAKENHI (JP17H04003 to K.H.), and AMED (20fk0108086j0202 to M.T.).

## Notes

### Competing Interest Statement

The authors have declared no competing interest.

## References

1. Nomoto, A., Koike, S. & Aoki, J. Tissue tropism and species specificity of poliovirus infection. Trends in microbiology 2, 47–51 (1994).

2. Shaghaghi, M. et al. New insights into physiopathology of immunodeficiency-associated vaccine-derived poliovirus infection; systematic review of over 5 decades of data. Vaccine 36, 1711–1719 (2018).

3. Macklin, G.R. et al. Evolving epidemiology of poliovirus serotype 2 following withdrawal of the serotype 2 oral poliovirus vaccine. Science 368, 401–405 (2020).

4. The World Health Organization, WHO Global Action Plan to minimize poliovirus facility-associated risk after type-specific eradication of wild polioviruses and sequential cessation of oral polio vaccine use. (WHO Press, Geneva, Switzerland; 2015).

5. Yasumura, Y. & Kawakita, Y. in Book, Studies on SV40 in tissue culture: preliminary step for cancer research in vitro. (eds. S. B. & T. Terasima) 1–19 (Department of Microbiology School of Medicine Chiba University, Chiba, Japan; 1988).

6. Osada, N. et al. The genome landscape of the African green monkey kidney-derived Vero cell line. DNA Res 21, 673–683 (2014).

7. Sakuma, C. et al. Novel endogenous simian retroviral integrations in Vero cells: implications for quality control of a human vaccine cell substrate. Sci Rep 8, 644 (2018).

8. Ellis, D.S. et al. Ebola and Marburg viruses: II. Thier development within Vero cells and the extra-cellular formation of branched and torus forms. J Med Virol 4, 213–225 (1979).

9. Ksiazek, T.G. et al. A novel coronavirus associated with severe acute respiratory syndrome. The New England journal of medicine 348, 1953–1966 (2003).

10. Drosten, C. et al. Identification of a novel coronavirus in patients with severe acute respiratory syndrome. The New England journal of medicine 348, 1967–1976 (2003).

11. Matsuyama, S. et al. Enhanced isolation of SARS-CoV-2 by TMPRSS2-expressing cells. Proceedings of the National Academy of Sciences of the United States of America 117, 7001–7003 (2020).

12. Zaki, A.M., van Boheemen, S., Bestebroer, T.M., Osterhaus, A.D. & Fouchier, R.A. Isolation of a novel coronavirus from a man with pneumonia in Saudi Arabia. The New England journal of medicine 367, 1814–1820 (2012).

13. Montagnon, B.J. & Vincent-Falquet, J.C. Experience with the Vero cell line. Dev Biol Stand 93, 119–123 (1998).

14. Barrett, P.N., Mundt, W., Kistner, O. & Howard, M.K. Vero cell platform in vaccine production: moving towards cell culture-based viral vaccines. Expert Rev. Vaccines 8, 607–618 (2009).

15. Montomoli, E. et al. Cell culture-derived influenza vaccines from Vero cells: a new horizon for vaccine production. Expert Rev. Vaccines 11, 587–594 (2012).

16. Warren, W.C. et al. The genome of the vervet (Chlorocebus aethiops sabaeus). Genome Res 25, 1921–1933 (2015).

17. Koike, S. et al. A second gene for the African green monkey poliovirus receptor that has no putative N-glycosylation site in the functional N-terminal immunoglobulin-like domain. J Virol 66, 7059–7066 (1992).

18. Ogawa, M. et al. Molecular mechanisms of Streptococcus pneumoniae-targeted autophagy via pneumolysin, Golgi-resident Rab41, and Nedd4-1-mediated K63-linked ubiquitination. Cellular microbiology 20, e12846 (2018).

19. Yamaji, T. et al. A CRISPR Screen Identifies LAPTM4A and TM9SF Proteins as Glycolipid-Regulating Factors. iScience 11, 409–424 (2019).

20. Manukyan, H. et al. Quantitative multiplex one-step RT-PCR assay for identification and quantitation of Sabin strains of poliovirus in clinical and environmental specimens. Journal of virological methods 259, 74–80 (2018).

21. Manukyan, H. et al. Multiplex PCR-based titration (MPBT) assay for determination of infectious titers of the three Sabin strains of live poliovirus vaccine. Virology journal 16, 122 (2019).

