## Supplementary Information for "Poliovirus-nonsusceptible Vero cell line for the World Health Organization global action plan"

**Supplementary Methods**

**Measles virus (MV) infection and virus titration.** The recombinant MV AIK-C strain expressing enhanced green fluorescent protein was used[1]. Vero and *PVR* KO cells seeded in a 12-well plate were infected with viruses at a MOI of 0.01. After 1 h incubation at 35°C, the cells were washed three times with PBS (−) and cultured at 35°C in a 5% $CO_2$ humid atmosphere with MEM containing 2% FBS and penicillin-streptomycin. At appropriate intervals, virus titer (plaque forming unit: PFU) in the culture fluid was determined by plaque assay[2]. Cells were dissociated by pipetting and collected together with culture supernatants as the culture fluid. Viral RNA copy number in the culture fluid was also determined by real-time RT-PCR, according to the Centers for Disease Control and Prevention (CDC) real-time measles RT-PCR protocol targeting the MV N gene (WHO: https://www.who.int/immunization/monitoring_surveillance/burden/laboratory/Annex_6.2.pdf?ua=1). For the real-time RT-PCR assay, RNA was extracted from 200 µl of the culture fluid with the High Pure Viral RNA kit (Roche, Mannheim, Germany) and 5 µl of RNA aliquot was used in each assay.

**Rubella virus (RV) infection and virus titration.** Vero and *PVR* KO cells seeded in a 12-well plate were infected with the recombinant RV expressing green fluorescent protein humanized monomer Azami Green 1 (AG1; rHS717AG1)[3] at MOI of 0.01. After 1 h incubation at 35°C, the cells were washed three times with PBS (−), and cultured at 35°C in a 5% $CO_2$ humid atmosphere with MEM containing 2% FCS and penicillin-streptomycin. Cells were dissociated by pipetting and collected together with culture supernatants as the culture fluid. At appropriate intervals virus titer in the culture fluid was determined by a focus forming assay[3]. Viral RNA copy number in the culture fluid was also determined by RV-specific real-time RT-PCR[4].

**Japanese encephalitis virus (JEV) infection and titration.** Nakayama strain of JEV[5] was a gift from Dr. Eiji Konishi (Research Institute for Microbial Diseases, Osaka University). Cells were seeded at a density of $3 \times 10^5$ cells/well in a 12-well plate (Corning Inc, N.Y., USA) one day before inoculation and then infected with JEV at a MOI of 0.01 at 37°C for 2 h. After the medium was replaced with the fresh normal culture medium, cells were cultured for four days. Extracellular infectious JEV particles were counted by plaque forming assay as described previously[5].

### Supplementary Figure legends

**Supplementary Figure 1.** Nucleotide and deduced amino acid sequence of Vero-*PVR1* and Vero-*PVR2*.

cDNAs encoding the full-size Vero-*PVR1* and Vero-*PVR2* were synthesized from Vero cell-derived total RNA and sequenced. Only one type of Vero-*PVR1* cDNA, of which sequences were identical to the previously reported AGM-*PVR1* sequence (which is deposited as *Cercopithecus aethiops* mRNA for poliovirus receptor AGM alpha 1 in the GenBank; see Supplementary Table 1), was obtained (**a**). Conversely, two types of Vero-*PVR2* cDNA sequences were obtained from the Vero cell genome (shown as "PVR2_1" and "PVR2_2") (**b**), presumably due to SNVs between the two alleles of the gene. Nucleotide sequences for the ORF with the initiator ATG (upper lines) and the deduced amino acid sequences (lower lines) are shown.

**Supplementary Figure 2.** Summary of mutant Vero cell lines in this study.

Information of indels found in the indicated cell lines are shown. Some chromatographs of DNA sequences showed mixed patterns, suggesting that different frame-shift mutations occurred in different alleles in Vero-*PVR1* and Vero-*PVR2*. The Vero Δ*PVR1*-1, Vero Δ*PVR2*-1, and Vero Δ*PVR1/2*-1 cell lines correspond to Vero Δ*PVR1,* Vero Δ*PVR2,* and Vero Δ*PVR1/2* cell lines, respectively, as described in the main text.

**Supplementary Figure 3.** Multiple rounds of PV replication in *PVR1* SKO, *PVR2* SKO, and *PVR1/PVR2* DKO cell lines at low or high MOI.

(**a**–**d**) Δ*PVR1*-1 (**a**, **b**), Δ*PVR2*-1 (**c**, **d**), and Δ*PVR1/2*-2 (**e**, **f**) were infected with PV Sabin 1 or 3 strains at a MOI of 0.01, and virus titers (**a**, **c**, **e**) and viral RNA (**b**, **d**, **f**) in the culture fluids (which was defined as a combined fraction of the retrieved cells and culture supernatant; see the main text) were quantified for 14 days. (**g**–**j**) Parental, Δ*PVR1/2*-2, and Δ*PVR2*-2 cells were infected with PV Sabin 1 (**g**, **h**) or 3 (**i**, **j**) strains at a MOI of 10, and virus titers (**g**, **i**) and viral RNA (**h**, **j**) were quantified up to seven days post infection (means ± S.D., $n$ = 3).

**Supplementary Figure 4.** Multiple rounds of measles virus (MV) replication in the parental, *PVR1* SKO, *PVR2* SKO, and *PVR1/PVR2* DKO cell lines.

Parental (**a**, **b**), Δ*PVR1*-1 (**c**, **d**), Δ*PVR2*-1 (**e**, **f**), and Δ*PVR1/2*-1 and Δ*PVR1/2*-2 (**g**, **h**) cells were infected with MV (AIK-C vaccine strain-derived recombinant virus) at a MOI of 0.01, and virus titers in the supernatants (**a**, **c**, **e**, **g**) and viral RNA (**b**, **d**, **f**, **h**) in the culture fluids were quantified at various intervals (means ± S.D., $n$ = 3).

**Supplementary Figure 5.** Multiple rounds of rubella virus (RV) replication in the parental, *PVR1* SKO, *PVR2* SKO, and *PVR1/PVR2* DKO cell lines.

Parental (**a**, **b**), Δ*PVR1*-1 (**c**, **d**), Δ*PVR2*-1, Δ*PVR2*-2 (**e**, **f**), and Δ*PVR1/2*-1 and Δ*PVR1/2*-2 (**g**, **h**) cells were infected with RV (rHS717AG1 wild strain-derived recombinant virus) at a MOI of 0.01, and virus titers (**a**, **c**, **e**, **d**) and viral RNA (**b**, **d**, **f**, **g**) in the culture fluids were quantified at various intervals (means ± S.D., $n$ = 3).

**Supplementary Figure 6.** Titration of infectious Japanese encephalitis virus (JEV) in the culture supernatant of parental, *PVR1* SKO, *PVR2* SKO, and *PVR1/PVR2* DKO cell lines.

(**a**–**d**) The indicated cells were infected with JEV (Nakayama strain) at a MOI of 0.01, and virus titers in the culture supernatant were quantified for four days post-infection (means ± S.D., $n$ = 3). For more information of the Vero mutant cell lines, see Supplementary Fig. 2.

### References


1. Seki, F., Someya, K., Komase, K. & Takeda, M. A chicken homologue of nectin-4 functions as a measles virus receptor. *Vaccine* **34**, 7–12 (2016).
2. Takeda, M. *et al*. Generation of Measles Virus with a Segmented RNA Genome. *J. Virol.* **80**, 4242–4248 (2006).
3. Sakata, M. *et al*. Short Self-Interacting N-Terminal Region of Rubella Virus Capsid Protein Is Essential for Cooperative Actions of Capsid and Nonstructural p150 Proteins. *J. Virol.* **88**, 11187–11198 (2014).
4. Okamoto, K., Fujii, K. & Komase, K. Development of a novel TaqMan real-time PCR assay for detecting rubella virus RNA. *J. Virol. Methods* **168**, 267–271 (2010).
5. Saito, K. *et al*. Comparative characterization of flavivirus production in two cell lines: Human hepatoma-derived Huh7.5.1-8 and African green monkey kidney-derived Vero. *PLoS One* **15** (4): e0232274. (2020).

a

```
   1 atggcccgaaccatggccgccgcatggcctccgctgctactgacgctactggagctgtcttggccacccccaggaaccggggacatcatcgtgcaggcgcccacccaggtgcccggcttc 120
   1  M  A  R  T  M  A  A  A  W  P  P  L  L  L  T  L  L  E  L  S  W  P  P  P  G  T  G  D  I  I  V  Q  A  P  T  Q  V  P  G  F  40

 121 ttgggcgactccgtgacgctgccctgctacctacaggtgcccggcatggaggagacacacgtgtcacagctgacttggtcacggcatggtgaatccggcagcatggccgtcttccaccaa 240
  41  L  G  D  S  V  T  L  P  C  Y  L  Q  V  P  G  M  E  E  T  H  V  S  Q  L  T  W  S  R  H  G  E  S  G  S  M  A  V  F  H  Q  80

 241 acgcagggccccaactattcggagcccaaacggctggaattcgtggccgccagactgggcacagagctgcgggatgcctcactgaggatgttcgggttgcgcgtcgaggatgaaggcaac 360
  81  T  Q  G  P  N  Y  S  E  P  K  R  L  E  F  V  A  A  R  L  G  T  E  L  R  D  A  S  L  R  M  F  G  L  R  V  E  D  E  G  N  120

 361 tacacctgcctgttcgtcacgttcccacagggcagcaggagcgtggatatctggctccgagtgcttgcgaagccccagaacacagctgaggttcagaaggtccagctcactggaaagccg 480
 121  Y  T  C  L  F  V  T  F  P  Q  G  S  R  S  V  D  I  W  L  R  V  L  A  K  P  Q  N  T  A  E  V  Q  K  V  Q  L  T  G  K  P  160

 481 gtgcccgtggcccgctgcgtctccacaggcggtcgcccgccggcccacatcacctggcactcagacctgggcgggatgcccaataccagccaggcgccagggttcctgtctggcacagtc 600
 161  V  P  V  A  R  C  V  S  T  G  G  R  P  P  A  H  I  T  W  H  S  D  L  G  G  M  P  N  T  S  Q  A  P  G  F  L  S  G  T  V  200

 601 actgtcaccagcctctggatttggtgccctcaagccaggtggacggcaagagtgtgacctgcaaggtggagcacgagagctttgagaagcctcagctgctgactgtgaacctcaccgtc 720
 201  T  V  T  S  L  W  I  L  V  P  S  S  Q  V  D  G  K  S  V  T  C  K  V  E  H  E  S  F  E  K  P  Q  L  L  T  V  N  L  T  V  240

 721 tactatcccccagaggtatccatctctggctatgataacaactggtacctcagccagaatgaggccaccctgacctgcgacgctcgcagcaacccagagcccacaggctacaactggagc 840
 241  Y  Y  P  P  E  V  S  I  S  G  Y  D  N  N  W  Y  L  S  Q  N  E  A  T  L  T  C  D  A  R  S  N  P  E  P  T  G  Y  N  W  S  280

 841 acgaccatgggtcccctgccacccttcgctgtggcccagggcgcccagctcctgatccgtcctgtggataaaccaatcaacacaactttcatctgcaatgtcaccaatgccctaggagct 960
 281  T  T  M  G  P  L  P  P  F  A  V  A  Q  G  A  Q  L  L  I  R  P  V  D  K  P  I  N  T  T  F  I  C  N  V  T  N  A  L  G  A  320

 961 cgccaggcagaactgaccgtccaggtcaaagagggacctcccagtgagccctcaggcatgtccagtaacatcatcatcttcctgattctgggaatcgtgattcttctgaccctcctgggt 1080
 321  R  Q  A  E  L  T  V  Q  V  K  E  G  P  P  S  E  P  S  G  M  S  S  N  I  I  I  F  L  I  L  G  I  V  I  L  L  T  L  L  G  360

1081 atcggggtttatttctatcggtccagatgttcccgtgagttcctttggtgccatcatctgtctccctcgagtgaggagcatgccagcgcctcggctaatgggtatatctcctattcagat 1200
 361  I  G  V  Y  F  Y  R  S  R  C  S  R  E  F  L  W  C  H  H  L  S  P  S  S  E  E  H  A  S  A  S  A  N  G  Y  I  S  Y  S  D  400

1201 gtgagcagagaggccagctcttcccaggatccacagacagagggcacaagg 1251
 401  V  S  R  E  A  S  S  S  Q  D  P  Q  T  E  G  T  R  417
```

**Supplementary Figure 1.** Nucleotide and deduced amino acid sequence of Vero-*PVR1* and Vero-*PVR2*.

b

```
PVR2_1 |    1 atggccgccgcatggcctccgctgctgctggcgctactggcgctgtcctggccacccccaggaaccggggacatcgtcgtgcaggcgcccacccaggtgcccggcttcttgggcgactcc 120
       |    1  M  A  A  A  W  P  P  L  L  L  A  L  L  A  L  S  W  P  P  P  G  T  G  D  I  V  V  Q  A  P  T  Q  V  P  G  F  L  G  D  S  40
PVR2_2 |                                    a

  121 gtgacgctgccctgctacctacaggtgcccggcatggaggtgacgcatgtgtcacagctgacttggtcacggcatggtgaatccggcagcatggccgtcttccaccaaacgcagggctcc 240
   41  V  T  L  P  C  Y  L  Q  V  P  G  M  E  V  T  H  V  S  Q  L  T  W  S  R  H  G  E  S  G  S  M  A  V  F  H  Q  T  Q  G  S  80
                                               a   a  c
                                               E

  241 cactattcggagcccaaacggcctggaattcgtggccgccaggctgggcacggagctgcgggatgcctcattgaggatgttcgggttgcgcgtcgaggatgaaggcagctacacctgcgtg 360
   81  H  Y  S  E  P  K  R  L  E  F  V  A  A  R  L  G  T  E  L  R  D  A  S  L  R  M  F  G  L  R  V  E  D  E  G  S  Y  T  C  V  120
                          g
                           V

  361 ttcttcatcttcccgcagggcaaaaggagcgtggatatctggctccgagtgcttgccaagccacagaacacagctgaggttcagaaggtccagctcactggaaagccggtgcccgtggcc 480
  121  F  F  I  F  P  Q  G  K  R  S  V  D  I  W  L  R  V  L  A  K  P  Q  N  T  A  E  V  Q  K  V  Q  L  T  G  K  P  V  P  V  A  160

  481 cgctgcgtctccacaggggggtcgcccgccggcccacatcacctggcactcagacctgggcgggatgcccaataccagccaggcgccagggttcctgtctggcacagtcactgtcaccagc 600
  161  R  C  V  S  T  G  G  R  P  P  A  H  I  T  W  H  S  D  L  G  G  M  P  N  T  S  Q  A  P  G  F  L  S  G  T  V  T  V  T  S  200

  601 ctctggattttggtgccctcaagccaggtggacggcaagagtgtgacctgcaaggtggagcacgagagctttgagaagcctcagctgctgactgtgaacctcaccgtctactaccccccca 720
  201  L  W  I  L  V  P  S  S  Q  V  D  G  K  S  V  T  C  K  V  E  H  E  S  F  E  K  P  Q  L  L  T  V  N  L  T  V  Y  Y  P  P  240

  721 gaggtatccttctctggctatgataacaactggtacctcagccagaatgaggccaccctgacctgcgacgctcgcagcaacccagagcccacaggctacaactggagcacgaccatgggt 840
  241  E  V  S  F  S  G  Y  D  N  N  W  Y  L  S  Q  N  E  A  T  L  T  C  D  A  R  S  N  P  E  P  T  G  Y  N  W  S  T  T  M  G  280

  841 cccctgccacccttcgctgtggcccagggcgcccaactcctgatccgtcctgtggataaaccaatcaacacaactttcatctgcaatgtcaccaatgccctaggagctcgccaggcagaa 960
  281  P  L  P  P  F  A  V  A  Q  G  A  Q  L  L  I  R  P  V  D  K  P  I  N  T  T  F  I  C  N  V  T  N  A  L  G  A  R  Q  A  E  320

  961 ctgaccgtccaggtcaaagagcgacctcccagtgagccctcaggcatgtccagtaacatcatcatcttcctgattctgggaatcgtgattcttctgaccctcctggggatcggactttat 1080
  321  L  T  V  Q  V  K  E  R  P  P  S  E  P  S  G  M  S  S  N  I  I  I  F  L  I  L  G  I  V  I  L  L  T  L  L  G  I  G  L  Y  360

 1081 ttctatcagtccagatattcctgtaaggagcatgccagcgcctcggctaatgggtatatctcctattcagatgtgagcagagaggccagcttttgccaggatccacagacagagggcaca 1200
  361  F  Y  Q  S  R  Y  S  C  K  E  H  A  S  A  S  A  N  G  Y  I  S  Y  S  D  V  S  R  E  A  S  F  C  Q  D  P  Q  T  E  G  T  400

 1201 agg 1203
  401  R  401
```

**Supplementary Figure 1.** Nucleotide and deduced amino acid sequence of Vero-*PVR1* and Vero-*PVR2*.

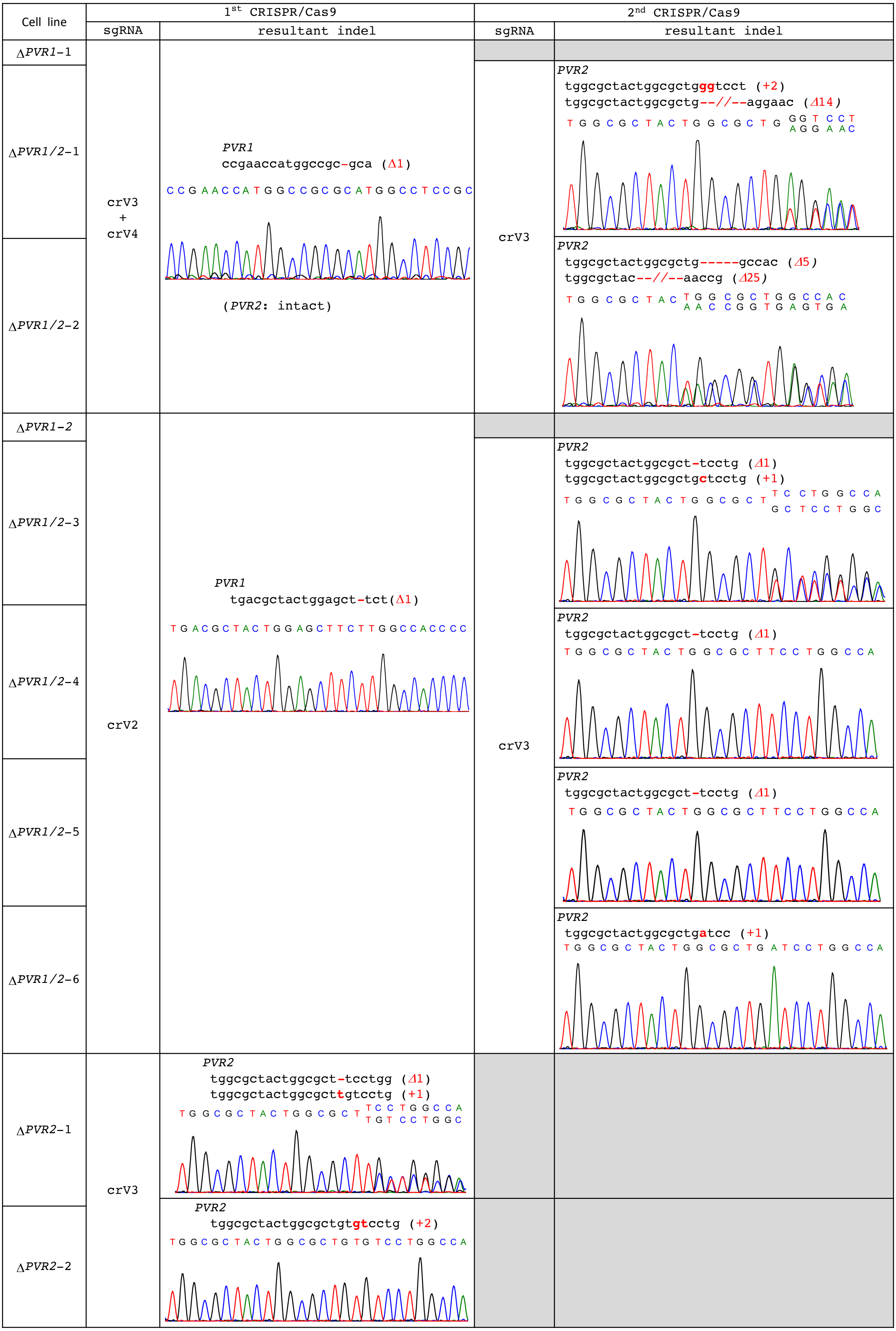


**Supplementary Figure 2.**
Summary of mutant Vero cell lines in this study.

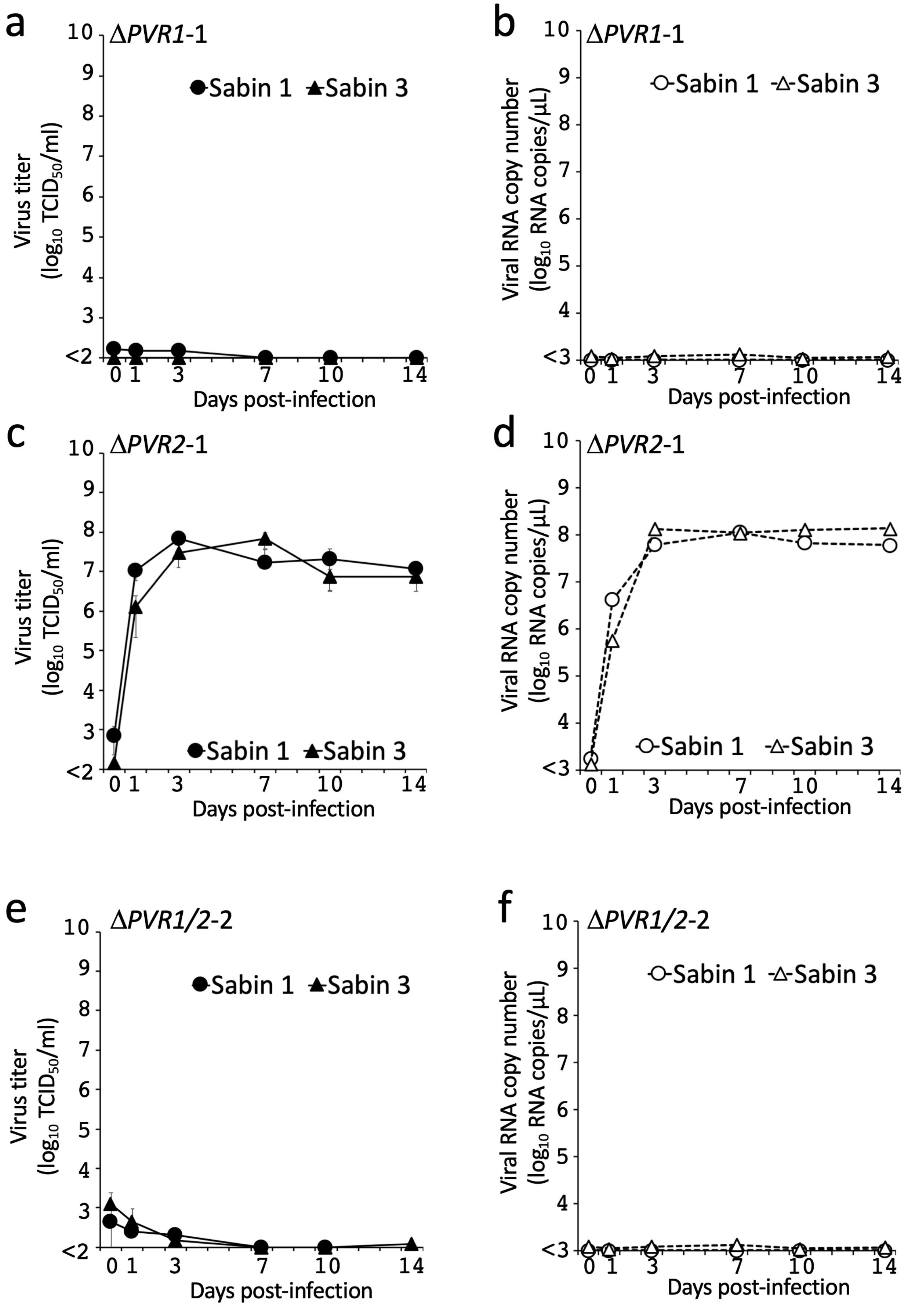


**Supplementary Figure 3.** Multiple rounds of PV replication in *PVR1* SKO, *PVR2* SKO, and *PVR1/PVR2* DKO cell lines at low or high MOI.

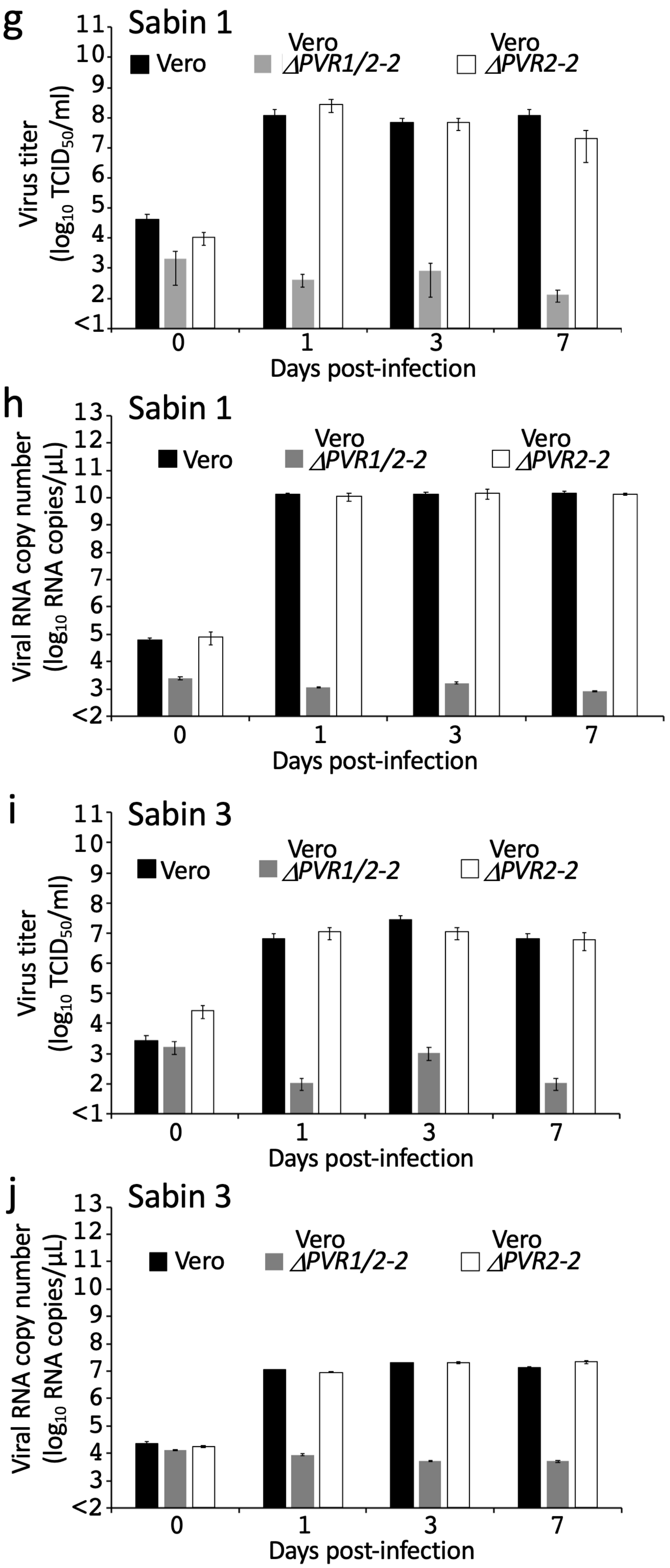


**Supplementary figure 3.** Multiple rounds of PV replication in *PVR1* SKO, *PVR2* SKO, and *PVR1/PVR2* DKO cell lines at low or high MOI.

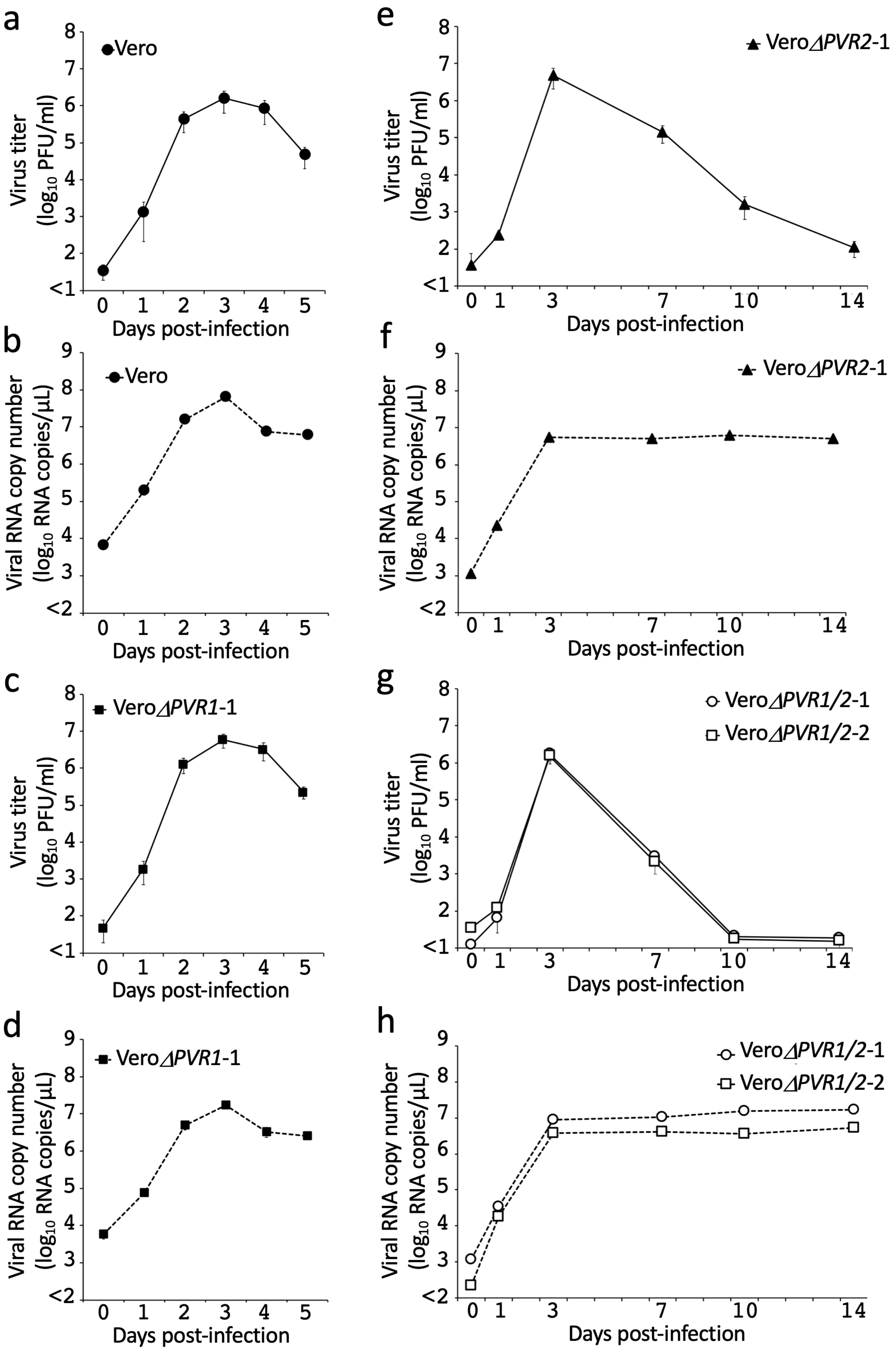


**Supplementary Figure 4.** Multiple rounds of measles virus (MV) replication in the parental, *PVR1* SKO, *PVR2* SKO, and *PVR1/PVR2* DKO cell lines.

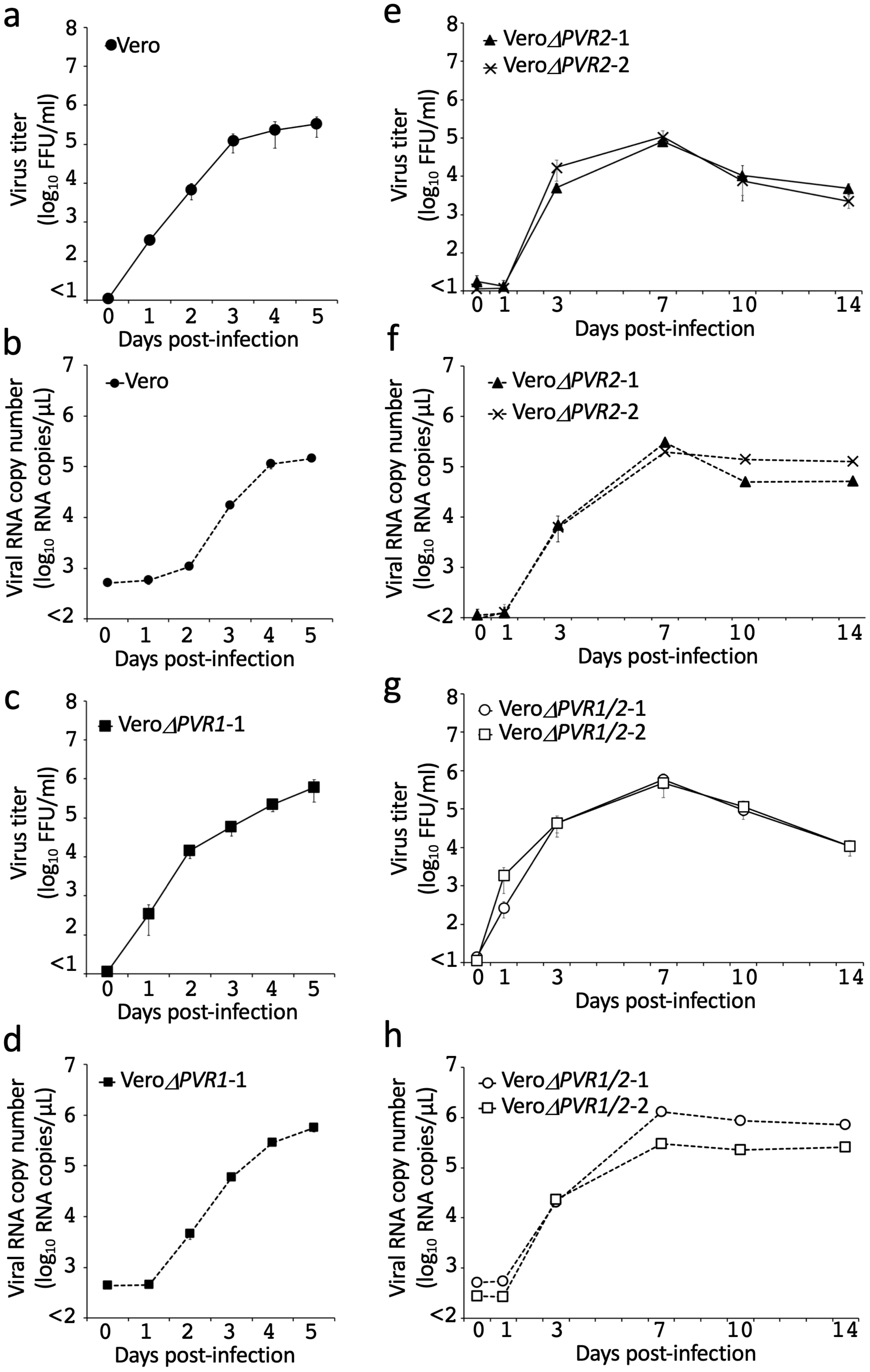


**Supplementary Figure 5.** Multiple rounds of rubella virus (RV) replication in the parental, *PVR1* SKO, *PVR2* SKO, and *PVR1/PVR2* DKO cell lines.

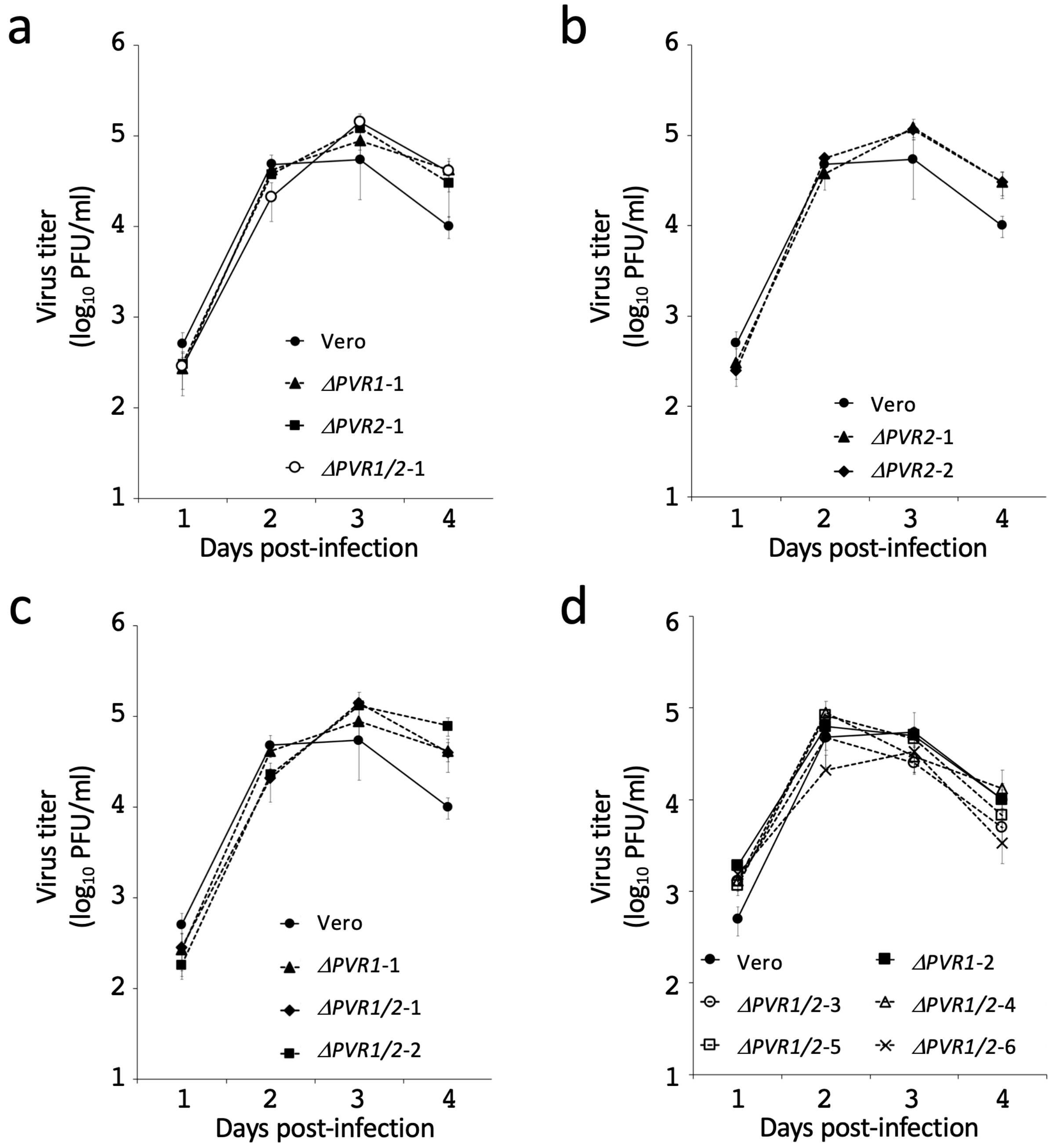


**Supplementary Figure 6.** Titration of infectious Japanese encephalitis virus (JEV) in the culture supernatant of parental, *PVR1* SKO, *PVR2* SKO, and *PVR1/PVR2* DKO cell lines.

**Supplementary Table 1. Database or Cell Resouce.**

| **Genomic, mRNA, or Protein Sequence** | |
|---|---|
| **Name** | **DATABASE** |
| Human poliovirus receptor (*PVR*), gene | Gene ID:5817 |
| Human *PVR*, protein | UniProtKB accession number: P15151 |
| African green monkey (AGM) *PVR1*, mRNA | GeneBank accession number: D12611.1 |
| AGM *PVR2*, mRNA | GeneBank accession number: D12613.1 |
| Vero *PVR1*, mRNA | This paper |
| Vero *PVR2*, mRNA | This paper |
| **Cell Lines** | **SOURCE** |
| Vero JCRB9013 (equivalent to the Vero ATCC CCL-81 cell line) ) | JCRB Cell Bank |
| Vero Δ*PVR1*-1, Δ*PVR1*-2, Δ*PVR2*-1, Δ*PVR2*-2, Δ*PVR1/2*-1, Δ*PVR1/2*-2, Δ*PVR1/2*-3, Δ*PVR1/2*-4, Δ*PVR1/2*-5, Δ*PVR1/2*-6 | This paper |
| HEp-2 | ATCC (CCL-23) |
